# Multiomics reveals the genomic, proteomic and metabolic influences of the histidyl dipeptides on heart

**DOI:** 10.1101/2021.08.10.455864

**Authors:** Keqing Yan, Zhanlong Mei, Jingjing Zhao, Md Aminul Islam Prodhan, Detlef Obal, Kartik Katragadda, Ben Doeling, David Hoetker, Dheeraj Kumar Posa, Liqing He, Xinmin Yin, Jasmit Shah, Jianmin Pan, Shesh Rai, Pawel Konrad Lorkiewicz, Xiang Zhang, Siqi Li, Aruni Bhatnagar, Shahid P. Baba

## Abstract

Histidyl dipeptides, are synthesized in the heart *via* enzyme carnosine synthase (Carns), which facilitates glycolysis and glucose oxidation by proton buffering and attenuate ischemia and reperfusion injury. However, a composite understanding of the histidyl dipeptide mediated responses in the heart are lacking. We performed multilayer omics in the cardio specific *Carns* overexpressing mice, showing higher myocardial levels of histidyl dipeptides lead to extensive changes in microRNAs that could target the expression of contractile proteins and enzymes involved in *β*-fatty acid oxidation and citric acid cycle (TCA). Similarly, global proteomics showed contractile function, fatty acid degradation and TCA cycle, pathways were enriched in the CarnsTg heart. Parallel with these changes, free fatty acids, and TCA intermediate-succinic acid were lower under aerobic and significantly attenuated under anaerobic conditions in the CarnsTg heart. Integration of multiomics data shows *β*-fatty acid oxidation and TCA cycle exhibit correlative changes at all three levels in CarnsTg heart, suggesting histidyl dipeptides are critical regulators of myocardial structure, function and energetics.

## Introduction

Glycolytically-active tissues-heart, skeletal muscle, and brain, contain small molecular weight histidyl dipeptides (227-241 Da), such as carnosine (*β*-alanine-histidine) and anserine (*β*-alanine-N^π^-histidine). These dipeptides are present at micro-milli molar levels in the myocardium that are synthesized by the ligation of a non-proteogenic amino-acid, β-alanine with histidine, *via* the enzyme carnosine synthase (Carns).^1, 2^ Because the pKa of histidyl dipeptides are close to the physiological pH (pKa 6.8-7.1), these dipeptides exhibit high buffering capacity, buffer intracellular pH and facilitate glycolysis during vigorous physical exercise and ischemia.^2, 3^ In addition to their high buffering capacity, histidyl dipeptides are efficient quenchers of reactive oxygen species, lipid peroxidation products, and they also chelate first transition metals.^4–7^ These dipeptides increase the expression of several metabolic enzymes, such as pyruvate dehydrogenase 4,^8, 9^ alter the release of microRNAs,^10^ and influence several signaling pathways, such as hypoxia inducible factor-α, ^11^ AKT/mTOR, ^12^ and STAT.^13^ In addition *β*-alanine, which is the rate limiting precursor amino acid for histidyl dipeptides,^14^ increases the expression of transcription factors, such as PPAR*δ*.^15^ The levels of histidyl dipeptides, within the heart and skeletal muscle can be increased by the dietary intake of β-alanine or by increased physical activity. In contrast their levels are decreased in the dysfunctional tissues such as failing heart.^14, 16^

Previous work shows perfusion of isolated mice and rat hearts with carnosine improves post ischemic contractile function.^5, 17^ Similarly, carnosine supplementation increases myocardial levels of carnosine and imparts protection against ischemia reperfusion (I/R) injury.^18^ Recently, we generated a cardio-specific Carns transgenic (CarnsTg) mice,^18^ which showed Carns overexpression significantly elevated the levels of histidyl dipeptides within the heart. Studying these mice, for the first time provided the opportunity to understand the effects of elevated myocardial histidyl dipeptides, independent of changes in physical activity and/or nutrition. In our work, we found that CarnsTg mice exhibit normal cardiac function, and increasing the myocardial production of histidyl dipeptides buffers intracellular pH, facilitates glucose utilization, and attenuates cardiac injury during I/R.^18^ Despite this evidence showing histidyl dipeptides imparts cardiac protection, improved glycolysis and influences numerous cellular and signaling pathways, an in-depth understanding of the histidyl dipeptide mediated effects on the genomic, proteomic and metabolomic landscape of the heart has not been performed. Thus, to dissect the effect of these dipeptides in the heart, we performed the genome-wide RNA sequencing (RNA-seq), global proteomics and untargeted metabolomics of the CarnsTg hearts. We integrated the three data sets to characterize the interactions, which are influenced by histidyl dipeptides and correlated at the gene, protein and metabolic levels. Further, we performed untargeted metabolomics in the CarnsTg heart after short durations of ischemia and determined the effects of these dipeptides on the cardiac fuel utilization in an ischemic heart.

Here we demonstrate that cardiac contraction, *β*-fatty acid oxidation and citric acid cycle (TCA) pathways were enriched at the protein levels in the CarnsTg heart. Parallel with these changes cardiac fuel utilization was improved by *Carns* overexpression under the aerobic and anaerobic conditions. Integration of the multi omics data set showed that TCA cycle and fatty acid degradation exhibit correlative changes at the microRNA, protein and metabolites levels in the CarnsTg heart. Hence this study is the first system level approach that combines the transcriptomics, proteomics and metabolomics data sets and provides new insights about the potential mechanisms by which histidyl dipeptides could improve cardiac metabolism and contraction, and thus alleviate cardiac injury associated with dysfunctional metabolic pathways and contractile dysfunction.

## Material and Methods

A detailed description of materials and methods is included in the Supplemental Methods.

### Statistical analysis

For metabolite identification, the maximum spectral similarity score is 1000 and the threshold of spectral similarity score was set as ≥ 500 (the maximum value is 1000). The p-value threshold set as *p* ≤ 0.001 for retention index matching. Partial least squares-discriminant analysis (PLS-DA), a supervised technique that uses the PLS algorithm to explain and predict the membership of samples to groups, was performed to give an overview on the metabolic profile difference among groups. A pairwise two-tail *t*-test with sample permutation was performed between the WT and the CarnsTg hearts, to determine whether a metabolite has a significant difference in abundance between the two groups. Grubbs’ test was employed for outlier detection before *t*-test. The *t*-test *p*-values were adjusted by up to 1000 times of sample permutation. The threshold of *t*-test was set *p* < 0.05. One-way Anova was performed to identify the differentially regulated metabolites in the WT and CarnsTg hearts subjected to perfusion only, perfusion followed by 5 and 15 min of ischemia. The data are normalized using the quantile method. All analysis were performed using SAS and R statistical software.

The RNA-Seq expression analysis was performed using NOISeq algorithm. ^19^ Transcripts with log2 fold-change *≥*2 and significant value > 0.8 were considered differentially expressed, as per the recommendation of the NOIseq. For proteins the fold change of *≥*2 and q-value <0.05 were considered as statistically significant. Pathway enrichment analysis of the differentially expressed proteins was performed based on GO and KEGG data base, and the statistical significance of the pathway were calculated using hypergeometric test. ^20, 21^

## Results

### Identification of differentially expressed genes in Carnosine synthase (Carns)-Tg heart

Carnosine synthase (Carns) participates in the synthesis of a wide range of histidyl dipeptides such as carnosine, anserine, and carcinine.^1^ To examine the effects of increasing the basal levels of this enzyme in the heart, we measured the levels of several histidyl dipeptides in the hearts of WT and CarnsTg mice. As we have reported previously^18^, the hearts of CarnsTg mice exhibit normal cardiac function, had significantly elevated levels of carnosine and anserine than WT hearts. In addition, in comparison with the WT, CarnsTg hearts had higher levels of homocarnosine as well, whereas no carcinine was detected (**Suppl. Fig. 1**). These observations suggest that the overexpression of *Carns* in the heart leads to a selective increase in carnosine, anserine and homocarnosine, but not other histidyl dipeptides.

Next to investigate whether elevated myocardial levels of histidyl dipeptides could affect the gene profile of the heart, we performed RNA-seq analysis of the WT and CarnsTg mice hearts and analyzed the gene expression changes. The paired-end sequencing of libraries generated 23.983 million sequencing reads, which after filtering for low quality generated 23.825 million clean reads. Of the total clean reads, the average mapping ratio of clean reads with reference gene was approximately 51.72%, 43.47% were unmapped reads and 4.81% were multi-position match. In comparison to 24,573 - the total number of mouse genes present in the database, we identified 17,509 genes in the WT and 17,468 genes in the CarnsTg mice hearts (**Data available on request**). Volcano plot of the RNA-seq data indicated the down-regulated (negative values) and the up-regulated (positive values) genes with log_2_ (fold change) *≥*2 and FDR value < 0.001 (**Fig. 1A**). We next profiled the differential gene expression comparing the WT and CarnsTg hearts, and identified a total of 100 differentially expressed genes (DEGs) in the CarnsTg heart of which 42 are the protein coding genes, and 21 of these protein coding genes were up-regulated and 21 genes were down-regulated. In the CarnsTg heart, the highly up-regulated genes were uncoupling protein 1, carbonic anhydrase 3, phosphoenolpyruvate carboxykinase 1, adiponectin, a predicted gene 38670 and histone cluster1 H2ag, and the significantly down-regulated genes included fos-like antigen2, RIKEN cDNA, ladybird homeobox homolog2 and SH3-binding domain kinase family, member 2. Among the 58-differentially expressed noncoding genes, 29 were up-regulated and 29 are down-regulated in the CarnsTg heart, which included the microRNA’s and small nucleolar RNAs. In the microRNA analyses, miR-7067, miR-142, miR-99a, miR-5625, miR-7036, and miR-5621 were highly up-regulated and miR-6989, miR-3101, miR-7091, and miR-5111 were significantly down-regulated (**Fig. 1B and Suppl. File I**).

**Figure 1:**
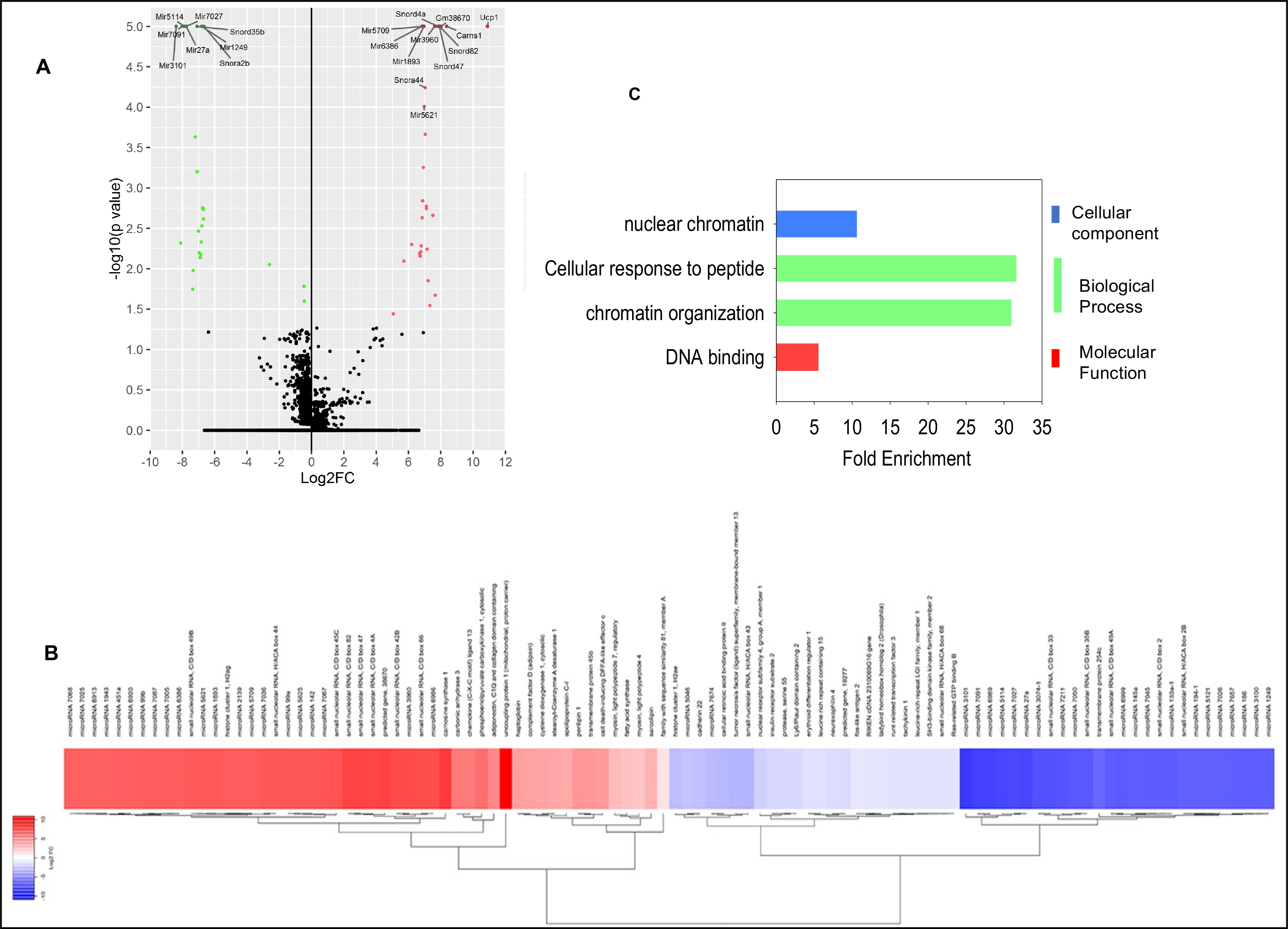
Transcriptomic analysis of the WT and CarnsTg hearts. (**A**) Volcano plot of the transcriptome between the CarnsTg and WT hearts. Statistical significance log_10_ of *p*-value Y-axis was plotted against log_2_-fold change (X-axis). (**B**) Heat map of the differentially expressed genes (DEGs) between the CarnsTg and WT hearts. (**C)** Gene ontology analysis of the differentially regulated non coding and coding genes between the CarnsTg and WT mice hearts, which were divided between three main categories: cellular component, biological component and molecular function.

To predict the biological processes linked to the altered gene expression, we functionally annotated all the 100 DEGs to the Gene Ontology (GO) enrichment analysis. For GO analysis, the annotated genes were divided into three major GO categories: cellular component, molecular function, and biological process. In the cellular component, the term of nuclear chromatin was the enriched component. Cellular responses to peptide and chromatin organization were the two top biological processes terms, and the most enriched component of the molecular function was DNA binding (**Fig. 1C**). Taken together these results suggests that increasing the myocardial levels of histidyl dipeptides affects the gene profile of heart.

### Proteomic profiling of the wild type (WT) and carnosine synthase (Carns) Tg hearts

While the flow of information from RNA to the protein translation is considered the central dogma, numerous studies have shown that a weak correlation exists between the mRNA expression and the abundance of translated proteins. ^24–28^ To determine whether the DEGs in CarnsTg heart could translate at the protein levels, we next examined the effect of elevated myocardial histidyl dipeptides on the protein profile and analyzed the differences in the protein expression between the WT and CarnsTg mice hearts by proteomic analysis. A total of 4719 and 4743 distinct proteins were identified from the WT and CarnsTg mice hearts, respectively (**Data available on request**). The differentially expressed proteins (DEPs) were identified with a q-value <0.01 and a fold change of *≥* between the WT and CarnsTg hearts. We found that approx. 939 proteins were differentially expressed between the WT and CarnsTg mice hearts (**Suppl. File II**). Of these, 566 were up-regulated and 373 proteins were down-regulated in the CarnsTg compared with the WT hearts. Principal component analysis of the DEPs clearly clustered the WT and CarnsTg proteins separately and unsupervised hierarchical clustering also resulted in grouping of WT and CarnsTg hearts into different clusters. The level of significance and magnitude of changes observed in the proteome by Carns overexpression was visualized by plotting the DEPs on a volcano plot (**Fig. 2A**). The highly up-regulated proteins in the CarnsTg compared with the WT hearts were the structural proteins such as troponin T, fast skeletal muscle, nebulin, myosin regulatory light chain 2, and titin. Furthermore, many key proteins that were up-regulated in the CarnsTg heart were the enzymes involved in the TCA cycle (isocitrate dehydrogenase, and succinate dehydrogenase), fatty acid transport and metabolism, such as carnitine palmitoyl transferase 2 (CPT2) and 2,3 enoyl-CoA hydratase. The significantly down-regulated proteins in the CarnsTg compared with the WT hearts were: aldehyde dehydrogenase, adipsin, and GTPase-activating protein.

**Figure 2:**
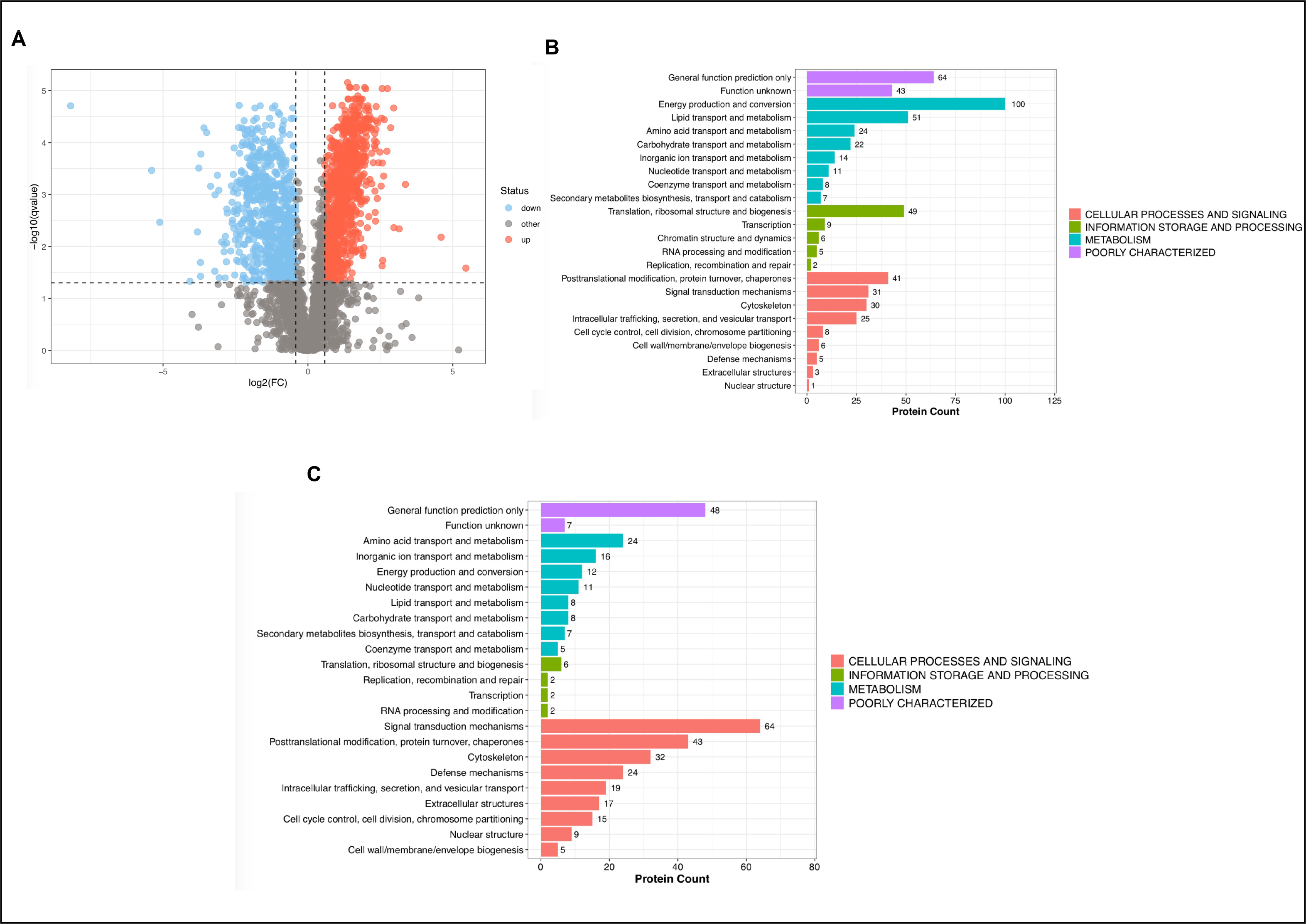
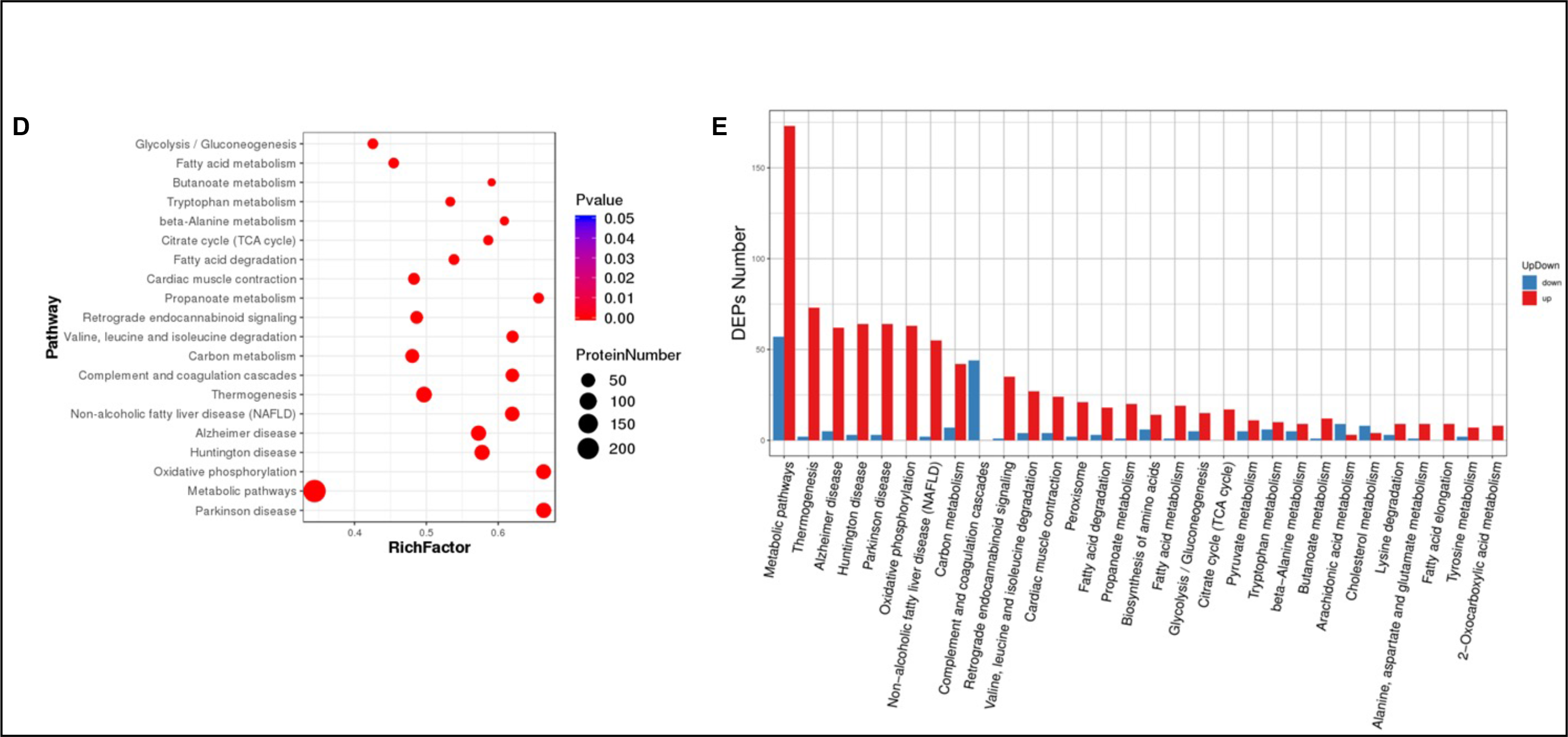
Comparative analysis, KOG and KEGG pathway analysis of the differentially expressed proteins (DEPs) between the WT and CarnsTg hearts. (**A**) Volcano plot of the proteome between the WT and Carns Tg hearts where the statistical significance log_10_ of *p*-value Y-axis was plotted against log_2_-fold change (X-axis). The Uniprot IDs of the total proteins identified in the WT and CarnsTg hearts via the MS/MS analysis were used to annotate the proteins with the corresponding KOG annotation. Functional enrichment analysis of, (**B**) upregulated showed that the greatest numbers of these proteins was allocated to metabolism and, (**C**) downregulated proteins showed the greatest number of these proteins are allocated to cellular processes and signaling. (**D**) KEGG enrichment analysis: the vertical axis represents the pathway terms with high enrichment and horizontal axis represents the Rich factor. The size of q-value is represented by the color of the dots. Smaller the q-value, the closer the color is towards red. (**E**) Enrichment of the specific KEGG pathway annotations for the upregulated (red) and the downregulated (blue) proteins in the CarnsTg hearts.

Next, we annotated the differentially expressed proteins to the NCBI annotation system gene ontology (GO) in order to identify the changes in specific biochemical pathways. The GO classification of the up-regulated proteins showed that the changes in cellular processes and signaling were enriched in the posttranslational modification, signal transduction and cytoskeleton. In the metabolism; energy production and conversion, lipid, amino acid, and carbohydrate transport and metabolism were highly enriched. In the information storage and processing; translation module-ribosome structure and biogenesis were highly enriched (**Fig. 2B**). In the down-regulated proteins; changes in the cellular processes and signaling were linked to signal transduction, post translational modification and cytoskeleton, and in metabolism changes in amino acid transport and metabolism were highly enriched (**Fig. 2C**). We then performed Kyoto Encyclopedia of genes and genomes (KEGG) enrichment analysis to analyze the DEPs, which are shown as scatter plot. KEGG pathway identified that DEPs in the CarnsTg mice heart were mostly enriched in metabolic pathways, glycolysis, oxidative phosphorylation, fatty acid degradation, cardiac muscle contraction and TCA cycle (**Fig. 2D**). Regarding the KEGG pathway functions associated with the up-regulated proteins, significant differences were found in the metabolic pathways, thermogenesis, oxidative phosphorylation, cardiac muscle contraction, fatty acid degradation and metabolism, glycolysis, TCA cycle and pyruvate metabolism. Pathway analysis of the down-regulated proteins in the CarnsTg heart were mainly enriched in the metabolic pathways, complement and coagulation cascades (**Fig. 2E**). Taken together, these results show Carns overexpression in the heart enriches the expression of enzymes involved in glycolysis, fatty acid oxidation and TCA cycle, and contractile proteins which could potentially influence the cardiac metabolism and contraction under diseased conditions.

### Effect of carnosine synthase overexpression on cardio metabolic profile under the basal and ischemic conditions

To investigate whether the changes observed in the expression of metabolic enzymes in the CarnsTg hearts could affect the cardiac metabolism, we compared the metabolic profiles of the WT and CarnsTg hearts under the basal and ischemic conditions. For this, polar metabolites from the snap-frozen hearts were extracted and derivatized with MTBSTFA and MSTFA and analyzed, using an unbiased and untargeted global metabolomics approach by GC*×*GC-MS. Approximately 2700 chromatographic peaks were detected in each of the MTBSTFA derivatized samples and 3700 chromatographic peaks were detected in the MSTFA derivatized samples. Approximately 280 and 520 metabolites were identified from the MTBSTFA and MSTFA derivatized samples respectively, including the fatty acids, amino acids, carbohydrates, glycolytic and citric acid (TCA) cycle intermediates and purines (**Suppl. File III and IV**). Partial least square discriminant analysis (PLS-DA) of the identified metabolites produced a clear separation between the WT and CarnsTg hearts (**Fig. 3A and B**). Volcano plot analysis of the differentially regulated metabolites identified after MTBSTFA and MSTFA derivatizations showed that 16 metabolites were significantly different in the CarnsTg compared with the WT hearts. Approximately, 12 metabolites were decreased and 4 metabolites were increased, in the CarnsTg compared with the WT hearts (**Fig. 3C and D**). Significantly, long chain fatty acid dodecanal and short chain fatty acids, such as propanoic acid, as well as the TCA intermediate, succinic acid and malic acid were lower in the CarnsTg heart, when compared with the WT hearts (**Table I**), suggesting that increased levels of histidyl dipeptides significantly impacts both fatty acid metabolism and the TCA cycle.

**Figure 3:**
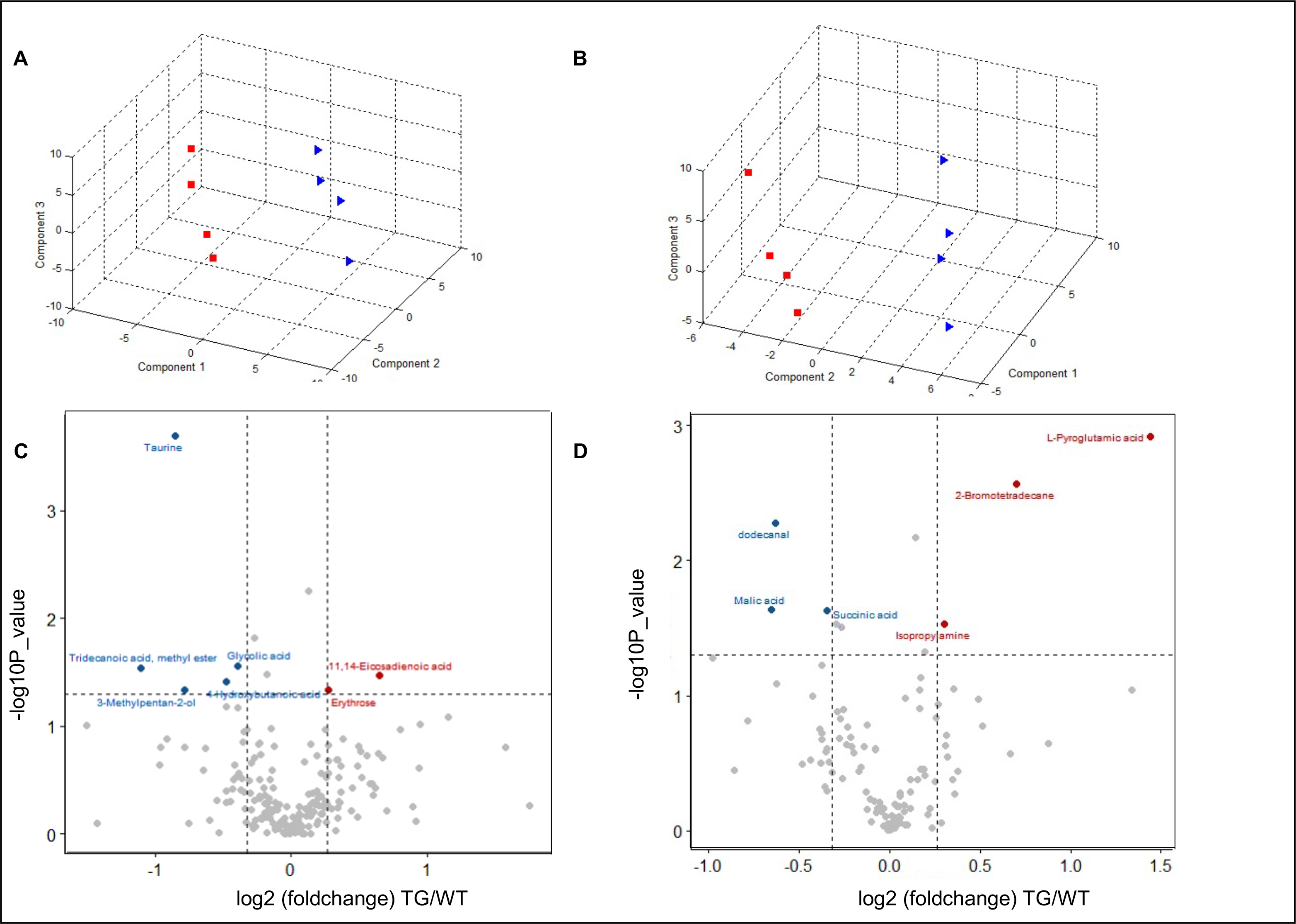
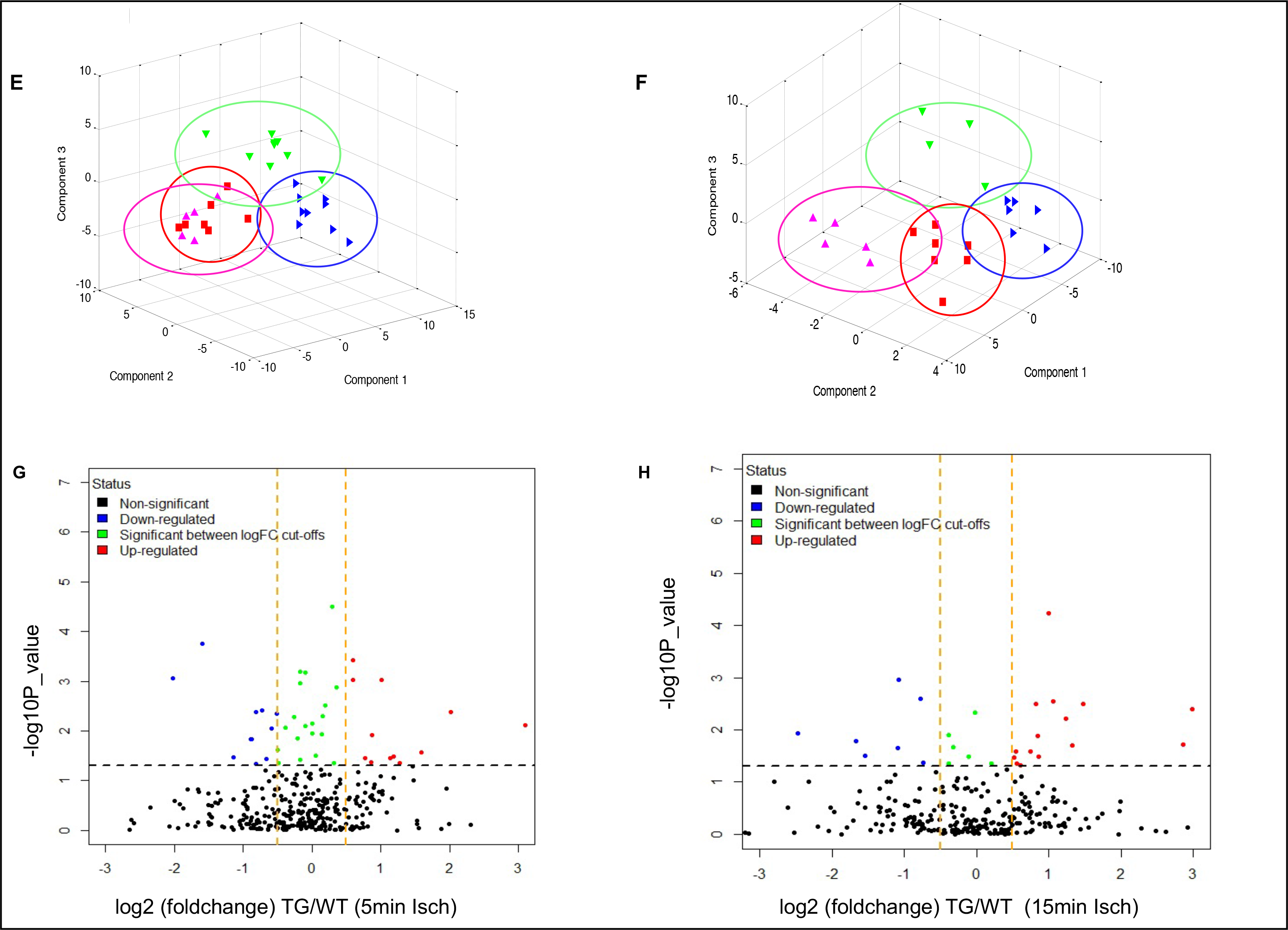
Metabolomic analysis of the WT and carnosine synthase (Carns) transgenic hearts under basal conditions and after short durations of ischemia. Changes in the global cardio metabolomic profile by Carns overexpression were assessed using an unbiased metabolomic approach. Partial least square plot (PLS-DA) of the metabolites from the WT and CarnsTg mice hearts under basal conditions detected by (**A**) MTBSTFA and (**B**) MSTFA derivatization. (**C, D**) Volcano plot represents the metabolites identified by MTBSTFA and MSTFA derivatization in the WT and Carns Tg hearts under the basal conditions. Isolated hearts from the WT and CarnsTg mice were subjected to 5 and 15 min of ischemia. Partial least square discriminant analysis (PLS-DA) of the WT and CarnsTg hearts following (**E**) 5 min and (**F**) 15 min of ischemia, where purple symbol represents metabolites from the WT heart subjected to 35 min perfusion, red symbol represents metabolites from CarnsTg heart perfused for 35 min, green symbols represents metabolites from WT heart subjected to 20 min perfusion followed by 5 min ischemia, blue symbols represents metabolites from CarnsTg heart perfused for 20 min followed by 15 min ischemia. Volcano plot represent the metabolites in the WT and Carns Tg hearts identified after (**G**) 5 min and (**H**) 15 min ischemia, n=5-7 mice hearts in each group.

**Table I.** Metabolites altered by *Carns* overexpression in the heart under basal conditions that were detected by MTBSTFA and MSTFA derivatization.

| No | Metabolites | p-value | Fold change | Change |
| --- | --- | --- | --- | --- |
| 1 | Taurine | 0.002 | 0.55 | Decrease |
| 2 | Amino butyric acid | 0.031 | 0.83 | Decrease |
| 3 | Carbonic acid | 0.05 | 0.51 | Decrease |
| 4 | Glycolic acid | 0.025 | 0.76 | Decrease |
| 5 | Erythrose | 0.032 | 1.16 | Increase |
| 6 | 4-Hydroxybutanoic acid | 0.038 | 0.72 | Decrease |
| 7 | 11,14-Eicosadienoic acid | 0.035 | 1.56 | Increase |
| 8 | 3-Methylpentan-2-ol | 0.045 | 0.58 | Decrease |
| 9 | dodecanal | 0.005 | 0.65 | Decrease |
| 10 | Catechol | 0.021 | 0.64 | Decrease |
| 11 | Succinic acid | 0.022 | 0.79 | Decrease |
| 12 | Isopropylamine | 0.027 | 1.23 | Increase |
| 13 | Pyroglutamic acid | 0.001 | 2.72 | Increase |
| 14 | Amino oxy acetic acid | 0.05 | 0.77 | Decrease |
| 15 | Propanoic acid | 0.028 | 0.81 | Decrease |
| 16 | Malic acid | 0.023 | 0.64 | Decrease |

Next to examine whether the increased myocardial levels of histidyl dipeptides affects the cardiometabolic profile during ischemia, which is independent of any neurohormonal effects, we subjected the isolated WT and CarnsTg hearts to different durations of global ischemia in the Langendorff mode. Hearts from the WT and CarnsTg mice were perfused for either (i) 35 min, (ii) 20 min followed by either 5 min and (iii) 15 min of global ischemia. The hearts were then immediately frozen for measuring the metabolites. Using an unbiased and untargeted global metabolomics approach, approximately 260 metabolites were identified. Analysis of the WT and CarnsTg perfused only (35 min) hearts showed that approx. 30 metabolites were different between the two groups (**Suppl. File V)**. PLS-DA of the metabolic profiles produced a clear separation between the WT and CarnsTg perfused hearts (**Suppl. Fig. 2A**). Volcano plot analysis showed that the abundance levels of several metabolites such as pyroglutamic acid, glycerol-3-phosphate, ribitol were increased, whereas several fatty acids, such as octadecanoic acid and arachidonic acid were decreased in the CarnsTg heart (**Suppl. Fig. 2B and File V**). To examine the influence of Carns overexpression on the global metabolomic profile in response to 5 min of global ischemia, we next created a PLS-DA plot with the samples classified into four groups: WT and CarnsTg hearts (perfusion only); WT and CarnsTg hearts (20 min perfusion and 5 min ischemia), which showed that the ischemia caused a significant deviation from the perfused hearts (**Fig. 3E**). Analysis of the metabolomic profiles between the WT perfused and ischemic hearts showed that approx. 60 metabolites were significantly different after 5 min of ischemia. Ischemia resulted in a significant reduction in amino acids proline and nor leucine), accumulation of free fatty acids (FFA; nonanoic acid and decanoic acid), and decreases in the metabolites of fatty acid (3-hydroxybutyric acid) and intermediates of TCA cycle (oxaloacetic acid; **Suppl. File V**). Metabolomic profiling between the WT and CarnsTg ischemic hearts (5 min) showed approx. 29 metabolites were significantly different between the two groups (**Table II; Suppl. File V**). Volcano plot analysis showed the levels of FFAs (decanoic acid, nonanoic acid and stearic acid ) were lower and pyruvic acid-the metabolite of glycolysis was increased in the CarnsTg compared with the WT ischemic hearts (**Fig. 3G**).

**Table II.**
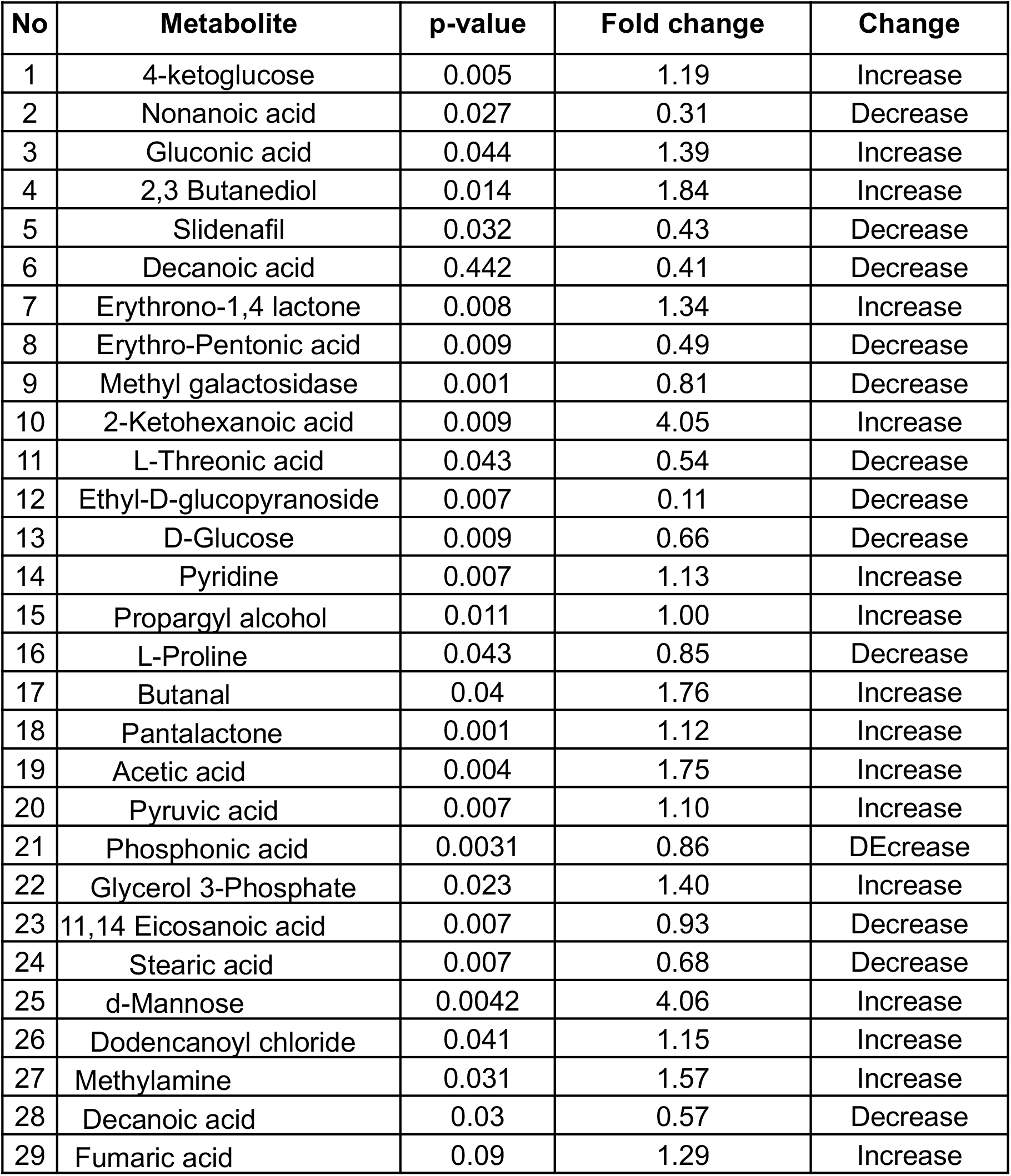
Metabolites significantly altered in the CarnsTg compared with the WT hearts after 5 mins of global ischemia.

| No | Metabolite | p-value | Fold change | Change |
| --- | --- | --- | --- | --- |
| 1 | 4-ketoglucose | 0.005 | 1.19 | Increase |
| 2 | Nonanoic acid | 0.027 | 0.31 | Decrease |
| 3 | Gluconic acid | 0.044 | 1.39 | Increase |
| 4 | 2,3 Butanediol | 0.014 | 1.84 | Increase |
| 5 | Slidenafil | 0.032 | 0.43 | Decrease |
| 6 | Decanoic acid | 0.442 | 0.41 | Decrease |
| 7 | Erythrono-1,4 lactone | 0.008 | 1.34 | Increase |
| 8 | Erythro-Pentonic acid | 0.009 | 0.49 | Decrease |
| 9 | Methyl galactosidase | 0.001 | 0.81 | Decrease |
| 10 | 2-Ketohexanoic acid | 0.009 | 4.05 | Increase |
| 11 | L-Threonic acid | 0.043 | 0.54 | Decrease |
| 12 | Ethyl-D-glucopyranoside | 0.007 | 0.11 | Decrease |
| 13 | D-Glucose | 0.009 | 0.66 | Decrease |
| 14 | Pyridine | 0.007 | 1.13 | Increase |
| 15 | Propargyl alcohol | 0.011 | 1.00 | Increase |
| 16 | L-Proline | 0.043 | 0.85 | Decrease |
| 17 | Butanal | 0.04 | 1.76 | Increase |
| 18 | Pantalactone | 0.001 | 1.12 | Increase |
| 19 | Acetic acid | 0.004 | 1.75 | Increase |
| 20 | Pyruvic acid | 0.007 | 1.10 | Increase |
| 21 | Phosphonic acid | 0.0031 | 0.86 | DEcrease |
| 22 | Glycerol 3-Phosphate | 0.023 | 1.40 | Increase |
| 23 | 11,14 Eicosanoic acid | 0.007 | 0.93 | Decrease |
| 24 | Stearic acid | 0.007 | 0.68 | Decrease |
| 25 | d-Mannose | 0.0042 | 4.06 | Increase |
| 26 | Dodencanoyl chloride | 0.041 | 1.15 | Increase |
| 27 | Methylamine | 0.031 | 1.57 | Increase |
| 28 | Decanoic acid | 0.03 | 0.57 | Decrease |
| 29 | Fumaric acid | 0.09 | 1.29 | Increase |

To identify the effects of histidyl dipeptides on the cardiometabolic profile, during longer durations of ischemia, we subjected the WT and CarnsTg hearts to either 35 or 20 min of perfusion, followed by 15 min of ischemia. PLS-DA plot of four groups: WT and CarnsTg hearts (perfusion only); WT and CarnsTg hearts (20 min perfusion and 15 min ischemia), produced a clear separation of the groups (**Fig. 3F**). We found that 63 metabolites were significantly altered in the WT ischemic heart, which belonged to the fatty acid, amino acid metabolism and TCA cycle (**Suppl. File V**). We also identified approx. 28 metabolites were significantly different between the WT *vs* CarnsTg ischemic hearts following 15 min of ischemia (**Suppl. File V, Table III**). Intermediates of glutathione cycle-pyroglutamic acid and TCA cycle (fumaric acid) were higher in the CarnsTg compared with the WT ischemic hearts (**Fig. 3H**). Levels of fatty acid (decanoic acid) and arachidonic acid were decreased and increased respectively in the ischemic CarnsTg heart. Taken together, these results suggests that the enrichment of enzymes involved in *β*-fatty acid oxidation and TCA cycle by *Carns* overexpression could optimize the cardiac fuel utilization under the basal and ischemic conditions.

**Table III.** Significantly altered metabolites in the CarnsTg compared with the WT hearts after 15 mins of ischemia.

| **No** | **Metabolite** | **p-value** | **Fold change** | **Change** |
| --- | --- | --- | --- | --- |
| 1 | 2-Pyrrolidinone | 0.011 | 2.104 | Increase |
| 2 | D-Mannose | 0.006 | 0.42 | Decrease |
| 3 | Decanoic acid | 0.042 | 0.6 | Decrease |
| 4 | 9-Octadecenoic acid | 0.0040 | 0.12 | Decrease |
| 5 | 4-Aminobutanoic acid | 0.011 | 5.54 | Increase |
| 6 | Fumaric acid | 0.031 | 2.91 | Increase |
| 7 | Mercaptoacetic acid | 0.032 | 0.55 | Decrease |
| 8 | D-glucose | 0.020 | 0.39 | Decrease |
| 9 | Ribitol | 0.020 | 1.25 | Increase |
| 10 | Phosphonic acid | 0.006 | 0.50 | Decrease |
| 11 | Pyroglutamic acid | 0.04 | 1.31 | Increase |
| 12 | Arachidonic acid | 0.04 | 1.31 | Increase |
| 13 | Pyruvic acid | 0.02 | 0.59 | Decrease |
| 14 | 2-Keto-hexanoic acid | 0.02 | 2.12 | Increase |

### Metabolic Pathway analysis

Given that Carns overexpression affects several different metabolic pathways, under both the basal and ischemic conditions, we next performed pathway analysis of the metabolites identified by MTBSTFA and MSTFA derivatization. This analysis showed that under the basal conditions, biochemical pathways involved in the metabolism of pentose phosphate pathway, valine, leucine and isoleucine, as well as arginine biosynthesis were different between the CarnsTg and WT hearts (**Fig. 4A and B**). Pathway analysis of the significantly different molecules, identified 2-3-fold enrichment in the TCA cycle, glycolysis, biosynthesis of unsaturated fatty acids, and 10-15-fold enrichment in the phenylalanine, tyrosine and tryptophan metabolism (**Fig. 4C and D**).

**Figure 4:**
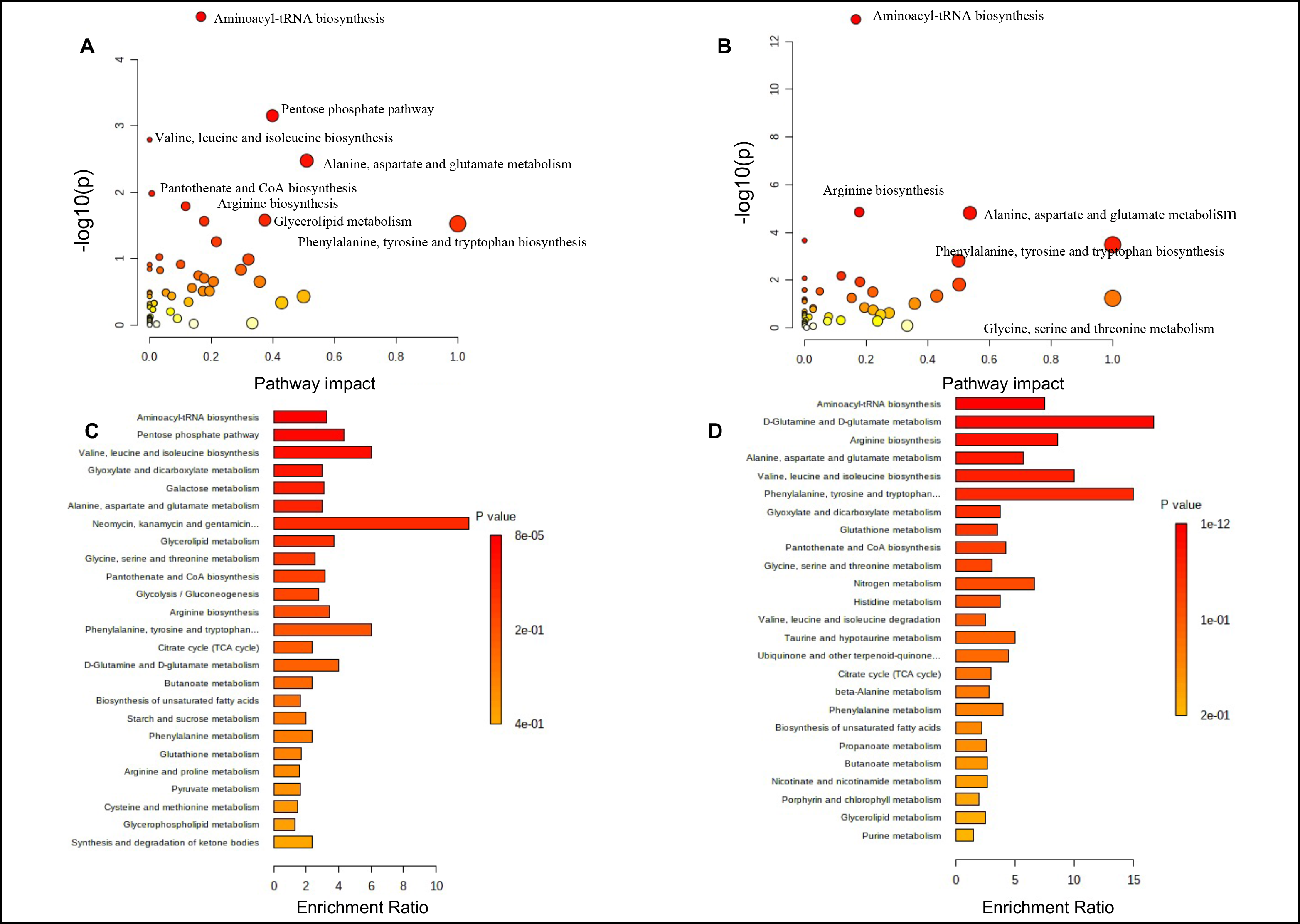
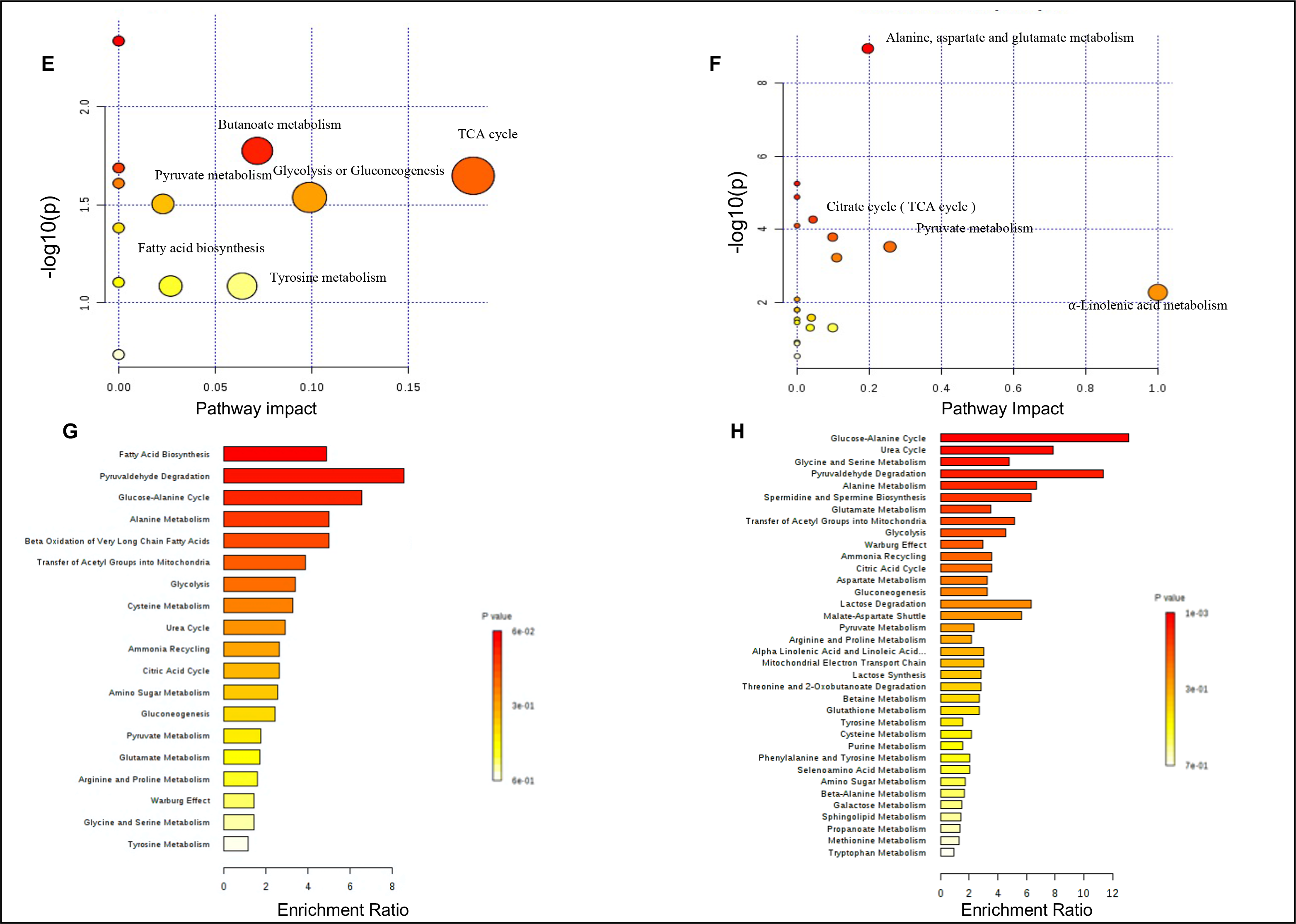
Pathway impact and enrichment of the differentially regulated metabolites in the WT and carnosine synthase (Carns) transgenic hearts. (**A and B**) Pathway impact analysis and the (**C and D**) pathway enrichment of the differentially regulated metabolites between the WT and CarnsTg mice hearts under basal conditions. Pathway impact analysis and pathway enrichments of the differentially regulated metabolites (**E and G**) after 5 min, ( **F and H**) and after 15 min of ischemia, n=5-7 hearts per group.

Next, we examined the metabolic pathways, affected by ischemia in the WT and CarnsTg hearts. Metabolic analysis revealed that 5 min of ischemia induced perturbations in the TCA cycle, fatty acids and amino acid metabolism (**Suppl. Fig. 3A**). Pathway enrichment analysis of the metabolites, which were significantly different, identified 4-10-fold enrichment in the malate-aspartate shuttle, urea cycle, and aspartate metabolism (**Suppl. Fig. 3B**). Similarly, prolonged ischemia of the WT mice heart (15 min) induced significant perturbations in the TCA cycle and amino acid metabolism, and significant enrichment of the malate-aspartate shuttle (**Suppl. Fig. 3C and D**). To examine the metabolic pathways, which are affected by Carns overexpression in the ischemic heart, pathway analysis of the metabolites that are different between the CarnsTg and WT mice hearts following 5 min of ischemia, indicated significant differences in glycolysis, TCA cycle, fatty acid biosynthesis, pyruvate, butanoate, arginine and proline metabolism (**Fig. 4E**). Enrichment analysis identified approximately 2-8-fold enrichment in pyruvaldehyde degradation, *β*-oxidation of very long chain fatty acids, alanine and cysteine metabolism, glucose-alanine cycle, glycolysis and TCA cycle (**Fig. 4G**). Furthermore, pathway impact analysis of metabolites which were differentially affected following 15 min of ischemia between the WT and CarnsTg hearts, showed that the highest pathway impact for the CarnsTg ischemic hearts was due to TCA cycle, alanine, aspartate and glutamate metabolism (**Fig. 4F**). Enrichment analysis of these metabolites identified 5-8-fold differences in the glucose-alanine cycle, pyruvaldehyde degradation, transfer of acetyl groups into mitochondria, TCA cycle and glycolysis between the WT and CarnsTg hearts (**Fig. 4H**). Taken together, these data suggest that increasing the myocardial levels of histidyl dipeptides improves multiple metabolic pathways under both the aerobic and anaerobic conditions, in particular the utilization of glucose and fatty acids.

### Integration of the transcriptomic, proteomic and metabolite networks

Given that the effects of elevating histidyl dipeptides were observed at the gene, protein and metabolite levels, we next integrated the transcriptomics, proteomic and metabolomic data sets and identified the levels of interaction between the three data sets. We first correlated the 42 DEGs and 810 DEPs and found that none of the DEGs and DEPs overlapped and correlated with each other. Since most of the DEGs did not correlate with the DEPs, we next performed the *in-silico* analysis between the highly regulated DEPs with the differentially expressed miRNAs using the TargetScan. We found that miR-5046, which was down-regulated in the CarnsTg heart could target the expression of Troponin T3 fast skeletal muscle and calsequestrin. Similarly, miR-6989 and miR-3100 that were decreased, could target the expression of succinate dehydrogenase and 2,3 enoyl-CoA hydratase respectively in the CarnsTg heart (**Table IV**). Further, miR-6913 that is increased was predicted to target aldehyde dehydrogenase protein expression.

**Table IV.** Target proteins of differentially regulated miRNA.

| miRNA | Change | Putative target gene | Protein | Change |
| --- | --- | --- | --- | --- |
| mmu-miR-5046 | Decrease | Tnnt3 | Troponin T3 fast skeletal muscle | Increase |
| mmu-miR-5046 | Decrease | Casq2 | Calsequestrin | Increase |
| mmu-miR-6913 | Increase | Aldh2 | Aldehyde dehydrogenase | Decrease |
| mmu-miR-6989 | Decrease | Sdha | Succinate dehydrogenase | Increase |
| mmu-miR-3100 | Decrease | Echs1 | 2,3 enoyl-CoA hydratase | Increase |

To uncover the potential interactions between the DEPs and metabolites in the CarnsTg hearts, we compared the proteomics and metabolomics clusters for enriched GO term and KEGG pathways. Based on the KEGG pathway analysis, the enrichment of *β*-fatty acid oxidation and TCA pathways at the protein levels were paralleled with the high enrichment of *β*-fatty acid oxidation and TCA cycle at the metabolite levels. The main enzymes regulating the abundance of free fatty acids (FFAs) are fatty acid synthase, long chain acyl CoA synthetase, CPT1 and 2, and four enzymes involved in *β*-fatty acid oxidation including acyl-CoA dehydrogenase, 2,3 enoyl CoA hydratase, 3 hydroxyacyl CoA dehydrogenase and 3-ketoacyl-CoA thiolase. Our proteomic screening identified all the transporters and enzymes involved in regulating fatty acids (**Suppl. File II**). We found that the expression of CPT2, and all the 4 enzymes involved in *β*-oxidation of fatty acids were higher in the CarnsTg than WT hearts (**Fig. 5 A-E**). In parallel with the increase in the expression of enzymes involved in fatty acid oxidation, global metabolomic profiling at baseline showed that the levels of FFAs, dodecanal and propanoic acid were decreased in the CarnsTg heart (**Table I**). Similarly, following 5 min of ischemia, decanoic acid and stearic were decreased in the CarnsTg heart (**Fig. 5 G and H**). Furthermore, the intermediates of TCA cycle, succinic acid and fumaric acid were decreased and increased respectively in the CarnsTg heart, under the basal conditions and following 15 min of ischemia respectively. Mirroring the changes in TCA cycle intermediates, proteomic analysis showed that the expression of enzyme succinate dehydrogenase, which oxidizes succinate to fumarate, was increased in the CarnsTg heart (**Fig. 6 A-D**). In addition, our integration analysis of the three data sets showed that the decrease in glycolic acid was mirrored by decreased expression of aldehyde dehydrogenase in the CarnsTg heart (**Fig. 6E and F**). To further connect the predicted putative upstream regulators that lead to altered proteome and metabolic changes in the CarnsTg hearts, integration of the KEGG pathway Database and TargetScan analysis showed that three miRNAs (miRNAs: 6989, 3100 and 6913) that could target the expression of three proteins (succinate dehydrogenase, 2,3 enoyl-CoA hydratase and aldehyde dehydrogenase) belonging to *β*-fatty acid oxidation, TCA cycle, and ethylene glycol pathways (**Fig. 5 and 6 Schemes**) are the synergistic networks that are correlated at all the three levels and by which histidyl dipeptides could optimize the cardiac fuel utilization under the aerobic and anaerobic conditions.

**Figure 5.**
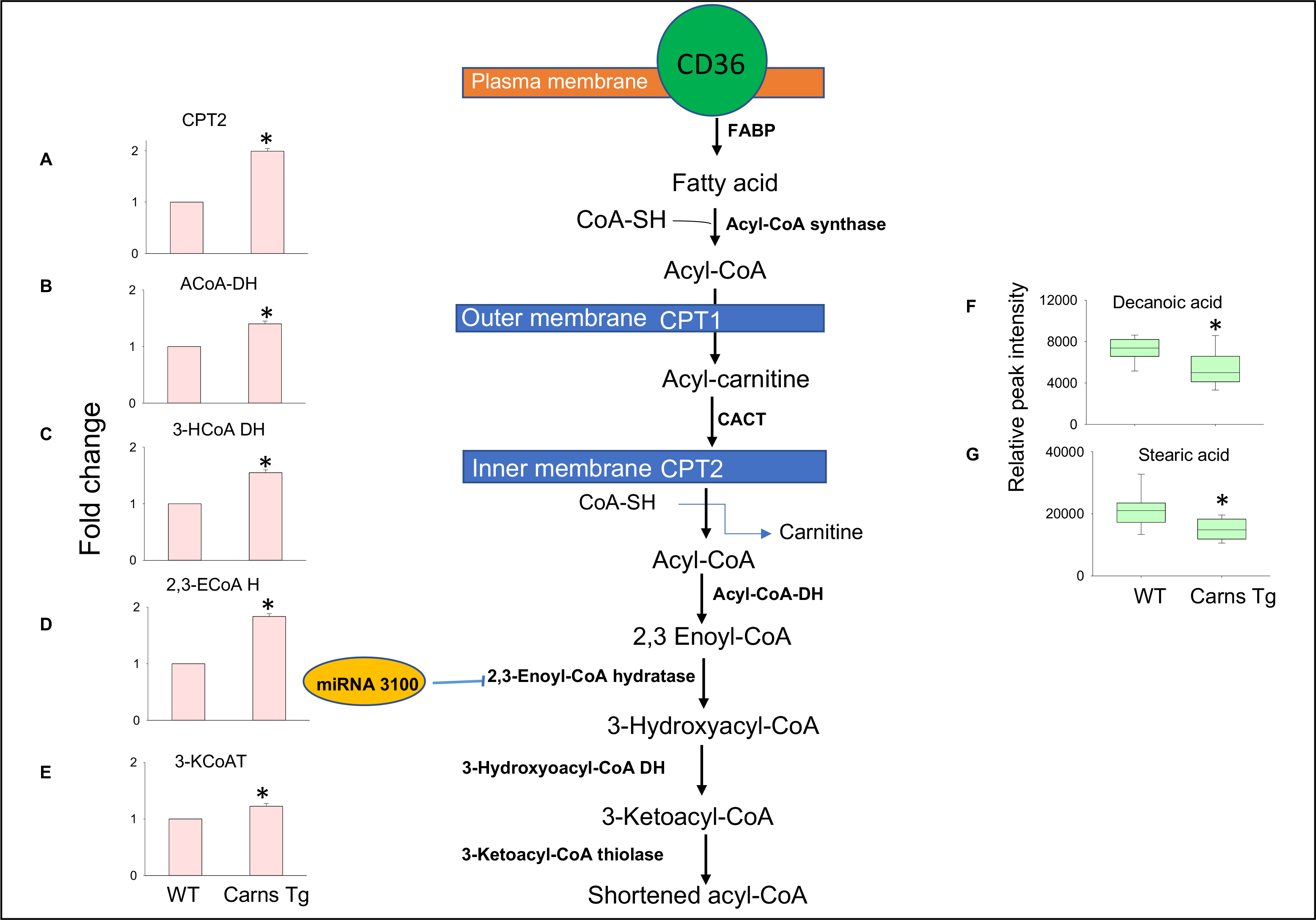
Transcriptomics, proteomic and metabolic interactions of the fatty acid metabolism in the CarnsTg heart. Schematic overview of the fatty acid metabolism, potential target of miRNA-3100 and levels of detected metabolites and proteins in the wild type (WT) and CarnsTg hearts. Relative fold changes in the expression of (**A**) carnitine palmitoyl transferase 2 (CPT2), (**B**) Acyl-CoA hydratase (ACoA-DH), (**C**) 3-hydroxyacyl-cCoA dehydrogenase (3HCoA DH) (**D**) 2,3 Enoyl-CoA hydratase (2,3-ECoAH), and (**F**) 3-Ketoacyl-CoA thiolase (KCoAT) between the WT and CarnsTg hearts. Free fatty acids levels (**G**) decanoic acid and (**H**) stearic acid in the WT and CarnsTg hearts following 5 min of global ischemia. *P<0.05 vs WT, n= 4-8 mice in each group.

**Figure 6.**
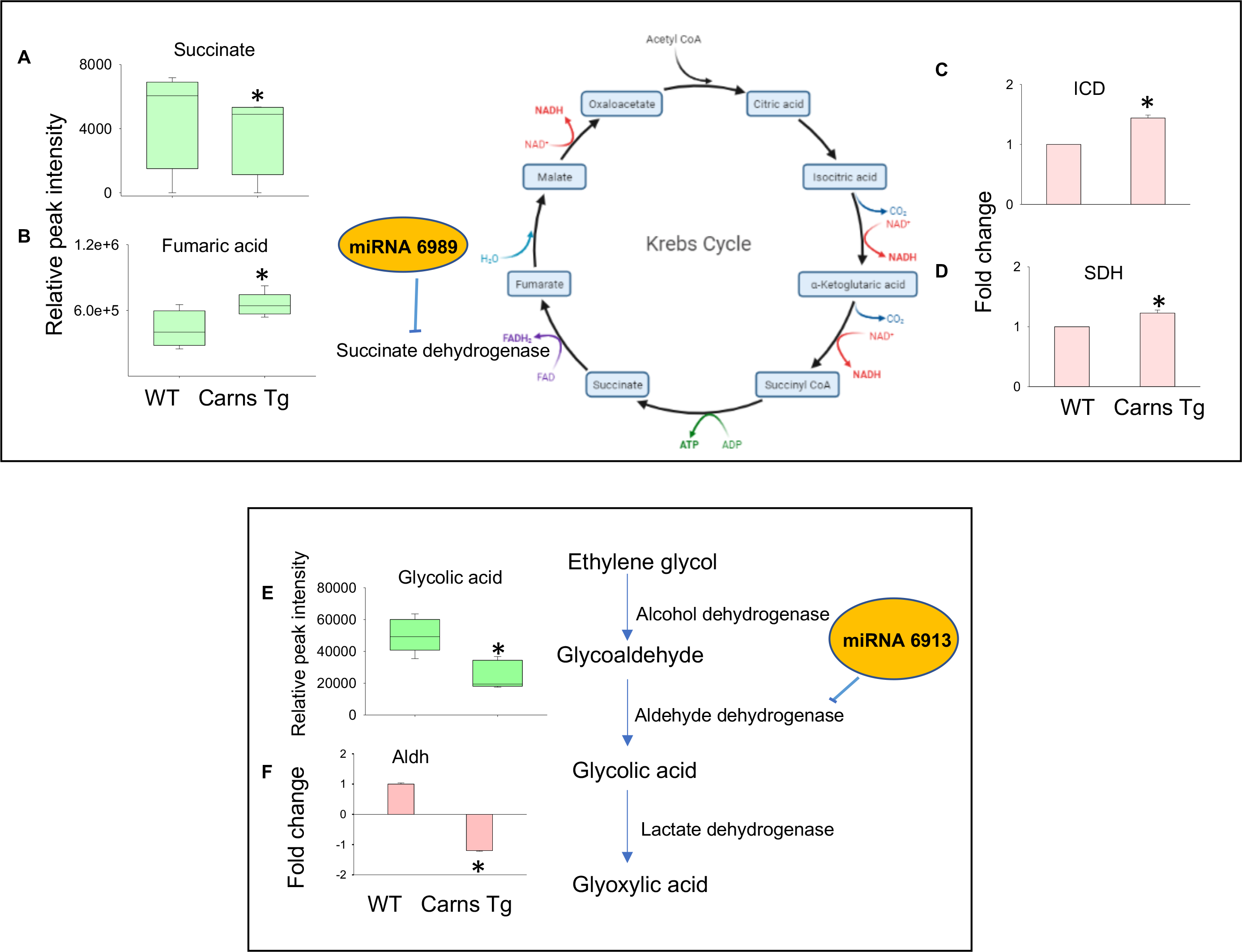
Transcriptomics, proteomic and genomic interactions in the CarnsTg hearts. Schematic overview of the citric acid cycle (TCA), potential target of miRNA-6989 succinate dehydrogenase and levels of TCA intermediates. (**A**) Levels of succinic acid under basal conditions and (**B**), fumaric acid following 15 min of ischemia. Relative fold changes in the expression of (**C**) isocitrate dehydrogenase (ICD) and (**D**) succinate dehydrogenase (SDH) between the WT and CarnsTg hearts. Schematic for glyoxylic acid formation and potential target of miRNA-6913 aldehyde dehydrogenase. Levels of (**E**) glycolic acid and (**F**) expression of aldehyde dehydrogenase in the WT and CarnsTg hearts, *p<0.05 vs WT, n= 4-8 mice in each group.

## Discussion

In the present study, we adopted a multi-layer omics approach to characterize in detail the genomic and proteomic changes occurring by increasing the myocardial histidyl dipeptide levels and then investigate their effects on the global metabolic profile, under the aerobic and anaerobic conditions. Further, we combined the RNAseq, global proteomics and untargeted metabolomics data sets to identify the metabolic pathways, which exhibited correlative changes on all the three levels. Transcriptomic analysis showed that the genes associated with chromatin organization and DNA binding, microRNAs that could target the expression of enzymes associated with β-fatty acid oxidation and citric acid cycle (TCA) were differentially regulated by histidyl dipeptides. Proteomic analysis showed that the abundance of approximately 939 proteins regulating different cellular, signaling and metabolic processes were affected by *Carns* expression. In particular, the abundance of several enzymes involved in the β-fatty acid oxidation and TCA cycle were enriched in the CarnsTg heart. Our metabolic profiling showed that the levels of free fatty acids were decreased and the intermediates of TCA cycle, such as fumaric acid was increased in the CarnsTg heart under the anaerobic conditions. Integration of the three data sets provided a comprehensive understanding of how the interactions at the miRNA and protein levels could possibly optimize the utilization of cardiac fuels, particularly fatty acids and glucose, under the physiological and pathological conditions in the CarnsTg heart. Collectively, our results for the first time provides an integrated resource of the genomic, proteomic and metabolomic changes that are affected by histidyl dipeptides in the heart. More broadly, results of concurrent genomic, proteomic and metabolic analyses highlight the limitation of these approaches individually, and illustrate how a combination of different approaches could provide a more comprehensive analysis of metabolic changes than by using each of these approaches individually.

### Role of histidyl dipeptides in gene and protein regulation.

Our omics analyses indicated that approximately 100 coding and non-coding genes and 938 proteins were differentially expressed in the CarnsTg heart and none of the differentially expressed genes (DEGs) and differentially expressed proteins (DEPs) were correlated at either the gene and protein levels. Among the most marked DEPs observed in the CarnsTg heart were troponin complex (I and T ), which increased 3-4-fold relative to the WT. It is interesting to note that *Carns* overexpression causes a switch and increases the expression of fast-skeletal muscle troponin in the heart. A number of regulatory networks such as thyroid hormone,^24^ transcription factors,^25^ and micro RNAs,^26^ could regulate the expression of the cardiac slow twitch and fast twitch contractile protein gene isoforms to the respective muscle type. Analysis of the data using the miRNA target prediction tool indicated that troponin T3 fast skeletal muscle is the possible target of miRNA-5046, suggesting that the down-regulation of miRNA-5046 observed in the CarnsTg could switch and increase the expression of fast-skeletal muscle, without discernable changes in the gene expression. Nevertheless, further investigations are required to confirm the presence of specific miRNA-protein interactions, and to garner more in-depth understanding of the mechanisms by which the post-transcriptional regulation of proteins is influenced by histidyl dipeptides.

One of the key findings of this work, not previously described is the enrichment of cardiac muscle contraction pathway in the Tg heart. The cardiac contractility is a complex machinery of cellular proteins, which includes tropomyosin *α*1 chain (TPM1), myosin light chain (MYL 1, 2, 3 and 4) and myosin heavy chain (MHC 6 and 7), and ATPase sarcoplasmic/endoplasmic reticulum. Our proteomic analysis showed that the expression of MYL1, 2 and 3, MHC7, troponin and calsequestrin were increased in the Tg hearts. A critical key to the pathogenesis of heart failure is the decreased expression of proteins involved in the contractile machinery, such as titin and myosin light chain.^27^ Previous reports show that MYL2 is decreased in the failing hearts ^28^ and mutation in the MYL2 lead to cardiomyopathy.^29^ Because contractile function of heart is the most important cardiac function, hence the upregulation of the sarcomere contractile machinery in the CarnsTg heart could possibly be a concerted mechanism to preserve the contractile dysfunction in the failing hearts.

### Role of histidyl dipeptides in basal myocardial metabolism

In our previous work we found that increasing the endogenous levels of histidyl peptides increases intracellular pH buffering. This increase in buffering capacity, allows the heart to maintain glycolysis during ischemia, generate higher levels of ATP and undergo less severe ischemic injury than hearts with lower buffering capacity.^18^ However, in addition to intracellular buffering, histidyl dipeptides can also affect other metabolic pathways. Previous studies show that carnosine induces the expression of pyruvate dehydrogenase,^8^ suggesting these ancillary changes may also be essential for preserving glycolysis or may have independent effects that may provide additional protection to the heart. Hence, to obtain a more holistic view of the metabolic effects of histidyl dipeptides, we performed an unbiased global metabolic profiling and found that 16 metabolites were significantly different in the CarnsTg mice hearts. We found that the levels of succinate, a TCA cycle intermediate, were lower in the CarnsTg than WT hearts. Concordant with this decrease, our proteomics analysis showed that in comparison with WT heart, the CarnsTg heart had increased expression of TCA cycle enzymes, particularly succinate dehydrogenase, which oxidizes succinate to fumarate. Succinate is recognized as a universal metabolic signature of ischemic injury that accumulates in the ischemic tissues upon reperfusion and drives a burst of reactive oxygen species (ROS) production by mitochondrial complex I. ^30^ Hence, the decrease of succinate in the CarnsTg heart under the basal conditions could potentially mitigate ROS production and ameliorate oxidative stress injury in the CarnsTg heart when subjected to ischemia, and thereby account for the better contractile recovery of the CarnsTg after ischemia.^18^

A key finding of this study is that the levels of long chain fatty acid dodecanal, as well as the levels of short chain fatty acids, such as propionic acid and 4-hydroxybutanoic acid were lower in the CarnsTg than WT heart. Fatty acids enter the cell *via* the fatty acid protein transporters, such as fatty acid translocase (CD36). ^31^ Then a family of fatty acid-binding proteins (FABP) facilitate fatty acid transport into cardiomyocytes.^32^ Free fatty acids are activated to form fatty acyl-CoAs, which are subsequently imported *via* the mitochondrial matrix by the carnitine shuttle system. Carnitine palmitoyl transferase I (CPT1) located in the outer membrane of mitochondria forms fatty acyl carnitine, which is translocated into mitochondrial membrane by carnitine-acylcarnitine translocase.^31^ Fatty acyl carnitine is converted back to fatty acyl CoA by carnitine-palmitoyl transferase II (CPT2) located on the mitochondrial membrane. ^31, 33^ In the mitochondria, *β*-oxidation of fatty acid is catalyzed by four enzymes acyl-CoA dehydrogenase, 2,3 enoyl CoA hydratase, 3-hydroxyacyl CoA dehydrogenase, and 3 ketoacyl-CoA thiolase. Although our proteomic analysis showed no change in the expression of CD36, the abundance of CPT2 and the 4 enzymes involved in the fatty acid oxidation was significantly higher in CarnsTg than WT hearts, suggesting that the enrichment of β-fatty acid oxidation in the CarnsTg heart observed at the proteome level, could potentially contribute towards decreasing the free fatty acid (FFA) levels.

The expression of fatty acid transporters and enzymes involved in β-fatty acid oxidation are largely under the transcriptional control of peroxisome proliferator-activated (PPAR)-*α* and *δ*, the retinoid X receptor-*α* and PPAR co-activator *γ* (PGC-1*α*).^34^ Previously, it has been reported that *β*-alanine, precursor for carnosine, increased PPAR*δ* expression suggesting that histidyl dipeptides could be the natural PPAR ligands.^15^ In addition, our *in-vivo* silico analysis showed that the miRNA-3100 decreased in the CarnsTg heart could target the expression of 2,3 enoyl-CoA hydratase. However, further studies are needed to confirm the mode of binding for these naturally occurring histidyl dipeptides with transcription factors and also validate whether the microRNAs affected by Carns overexpression could target the expression of enzymes involved in *β*-fatty acid oxidation. Nonetheless, our results so far indicate that multiple metabolic pathways are affected by enhancing myocardial histidyl dipeptides. Significantly in the CarnsTg heart, the decrease in free fatty acids is accompanied by an increase in the expression of the enzymes involved in β-fatty acid oxidation, suggests that higher levels of histidyl dipeptides are conducive to or permissive of fatty acid oxidation even in a non-stressed, non-ischemic heart.

### Role of histidyl dipeptides in the ischemic heart

Metabolic derangement is a key feature of cardiac ischemic injury. Multiple studies have shown that progressive decrease of glucose and fatty acid utilization is a characteristic signature of the ischemic hearts. ^33, 35–37^ Elevated levels of circulating FFAs in the ischemic heart are associated with increased incidence of ventricular arrhythmias, post infarction angina, and death in patients with acute myocardial infarction. ^33, 38, 39^ High levels of FFAs are also common in patients suffering from myocardial ischemia. Studies in animal models of regional and global ischemia show that excess fatty acid accumulation in the heart impairs ventricular function. Such accumulation of FFAs during ischemia, particularly long chain fatty acyl-CoA and long-chain acylcarnitine esters, weakens the membrane and compromises the function of membrane bound proteins. It has also been associated with increased mitochondrial membrane permeability, changes in calcium homeostasis, suppression of glucose utilization, and increased myocardial oxygen consumption in ischemic hearts. ^40–42^ In view of these findings, it has been suggested that decreasing the uptake and utilization of the fatty acids could be a potential strategy to salvage the ischemic myocardium. Although inhibiting the mitochondrial uptake of fatty acid by inhibiting CPT1 improves the post ischemic contractile function, ^43–, 45^ which is independent of decreasing fatty acid utilization, ^46^ decreasing the fatty acid oxidation under conditions of elevated fatty acid availability leads to the accumulation of toxic lipid intermediates including diacylglycerol, which cause aberrant cardiac signaling and toxicity. ^31, 47–51^ In the present study, we found that FFAs (nonanoic acid and decanoic acid), were increased in the isolated WT ischemic hearts, suggesting that the accumulation of FFAs is independent of their uptake and transport. We also found that Carns overexpression increased the utilization of FFAs and glucose, which was marked by decreases in the levels of stearic and decanoic acid and an increase in the pyruvic acid levels. This favorable profile could in turn attenuate ischemic injury, by preventing FFA-induced changes in cardiac signaling, metabolism, and mitochondrial function.

We had previously reported that *Carns* overexpression delays myocardial acidification during ischemia, which facilitates glucose metabolism via glycolysis.^18^ In the current study, our proteomic analysis showed that Carns overexpression increases the expression of the fatty acid transporters and enzymes involved in fatty acid metabolism. Therefore, the improvements in the fatty acid utilization observed under the ischemic conditions in the CarnsTg ischemic hearts could be either secondary to the improvements in pH and glucose utilization or due to the increase in the expression of β-fatty acid oxidation enzymes. In addition to free fatty acid accumulation, ischemia also induces substantial changes in the TCA cycle intermediates. It has been reported that the loading of TCA intermediates, such as fumarate, protects the heart from I/R injury. In studies with the mice deficient in fumarate dehydratase increases in cardiac fumarate levels are associated with marked reduction in infarct size.^52^ Our results show that fumaric acid was increased in the ischemic CarnsTg heart, which was mirrored by the increased expression of the succinate dehydrogenase that catalyzes the oxidation of succinate into fumarate. Taken together, the results of our metabolomic and proteomic profiling for the first time show that increasing the myocardial histidyl dipeptides enriches β-fatty acid oxidation pathway at the protein level, which is accompanied by increase in fatty acid and glucose utilization. Collectively these findings suggest that during ischemia, the elevated levels of histidyl dipeptides could support the channeling of both the cardiac fuels, glucose and fatty acids, through the TCA cycle.

### Multi-omics approach elucidates the impact of histidyl dipeptides

Overall, by using a well-defined model of a single gene overexpression, localized to the heart, we were in this study able to identify multiple transcriptomic, proteomic and metabolomic changes influenced by histidyl dipeptides in the heart. Many of the proteomic changes were concordant with the changes in metabolism and the overall metabolic protection provided by histidyl dipeptides to the ischemic heart. During the course of this work, we identified the previously unknown metabolites, transcripts and proteins, which were affected by histidyl dipeptides in the heart, and how these changes act in a concert to create a metabolic milieu in the heart that resists ischemic injury. To our knowledge, this is the first study using the system biology approach, to identify the metabolic, molecular and cellular pathways in ischemic and non-ischemic hearts and how they are affected by increasing the synthesis of histidyl dipeptides in the heart. Of importance, multi-omics integration of the data revealed potential parallels at the genome, proteome and metabolome levels in the heart, showing that the well-established fatty acid metabolism and TCA cycle, enriched at the microRNA and protein levels, overlap with the enrichment of fatty acid oxidation and TCA cycle metabolites. The unbiased assessment of the metabolic changes in the ischemic heart, and the findings that ischemic accumulation of FFA was attenuated in the CarnsTg hearts reveals novel insights into the reprogramming at the gene and protein levels, which impacts both β-fatty acid oxidation and glycolysis. Although we could not identify the individual contribution of carnosine, anserine, and homocysteine, our results suggest that a combinatorial effect of these dipeptides on multiple pathways, particularly fatty acid and glucose utilization could potentially contribute towards optimizing the use of cardiac fuels under the physiological and pathological conditions to impart a pervasive protection against ischemic injury. Given that the histidyl dipeptides in the heart are increased by exercise,^18^ our findings provide a new understanding of the mechanisms underlying the cardioprotective effects of exercise. Finally, because myocardial levels of histidyl dipeptides could be readily increased by feeding carnosine or *β*-alanine, these results also suggest a readily adaptable intervention to improve basal metabolism and contractile function in the heart and to increase myocardial resistance to ischemic injury.

#### Authors contributions

S.P.B., A.B., and S.L conceptualized and designed the study. KY, ZM, J.Z., contributed equally to sample collection, sample processing, data acquisition, analysis and interpretation. M.A.I.P., D.O., K.K., L.H., J.C., D.P.K., L.H., X.Y., J.S., P.K.L., X.Z., S.R., J.P., and B.D. were responsible for sample processing, data acquisition and analysis. All authors reviewed and contributed to the manuscript.

## Acknowledgments

We thank D. Mosley, F. Li and the Diabetes and Obesity Center’s Imaging and Physiological Core and Animal Models and Phenotyping Core at the University of Louisville for technical assistance.

## Sources of Funding

This work was supported by the National Institutes of Health: R01HL122581-01 (SPB), R01HL 55477, GM127607 (AB).

## Conflict of Interest Disclosures

None

## Footnotes

*KY, ZM, J.Z. contributed equally

---

**Supplemental Fig. 1.**
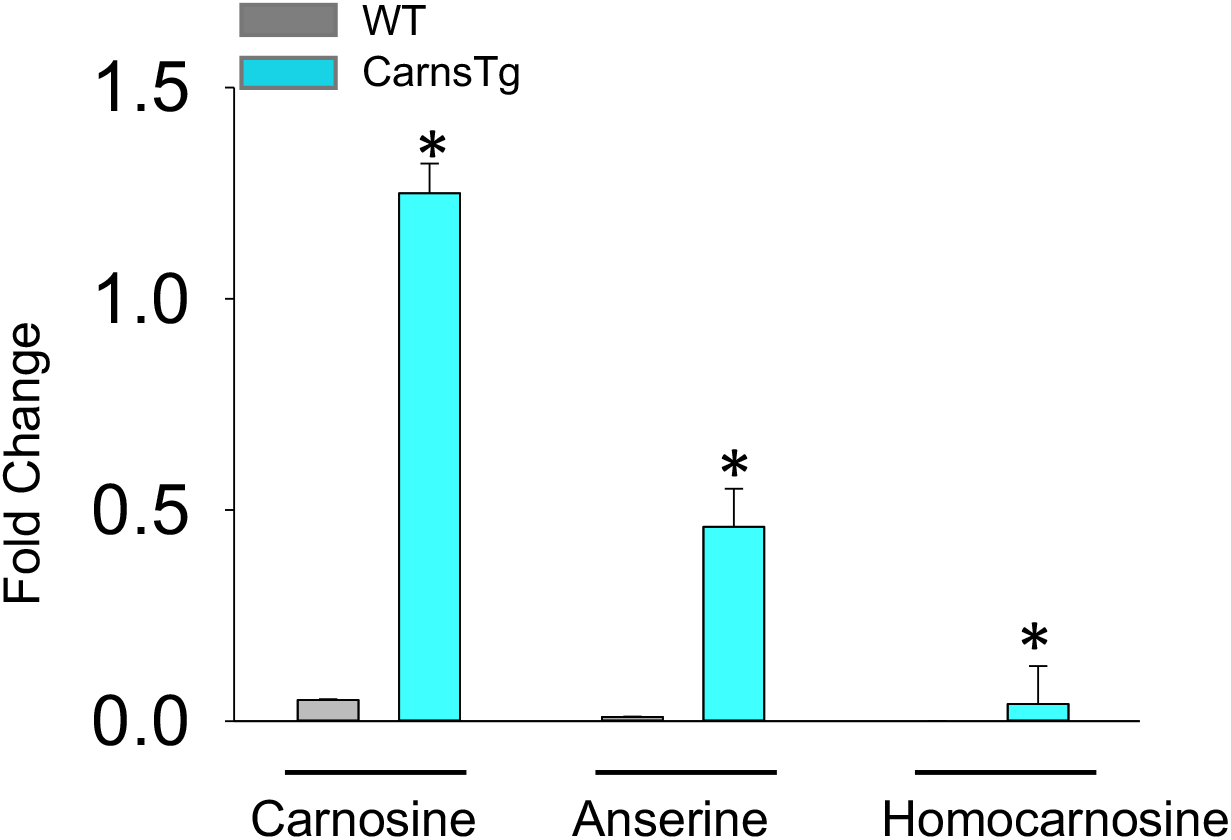

**Supplemental Fig. 2.**
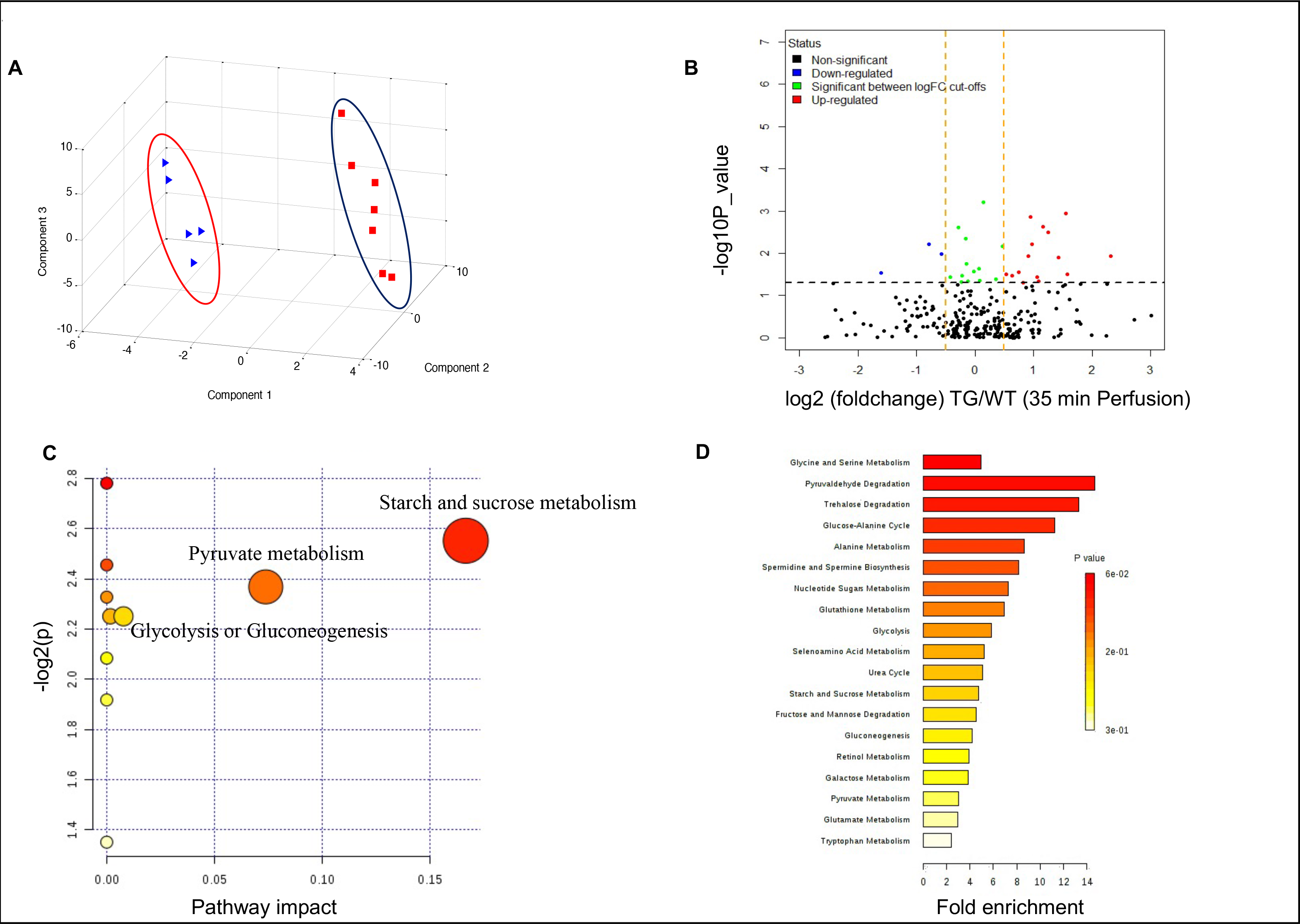

**Supplemental Fig. 3.**
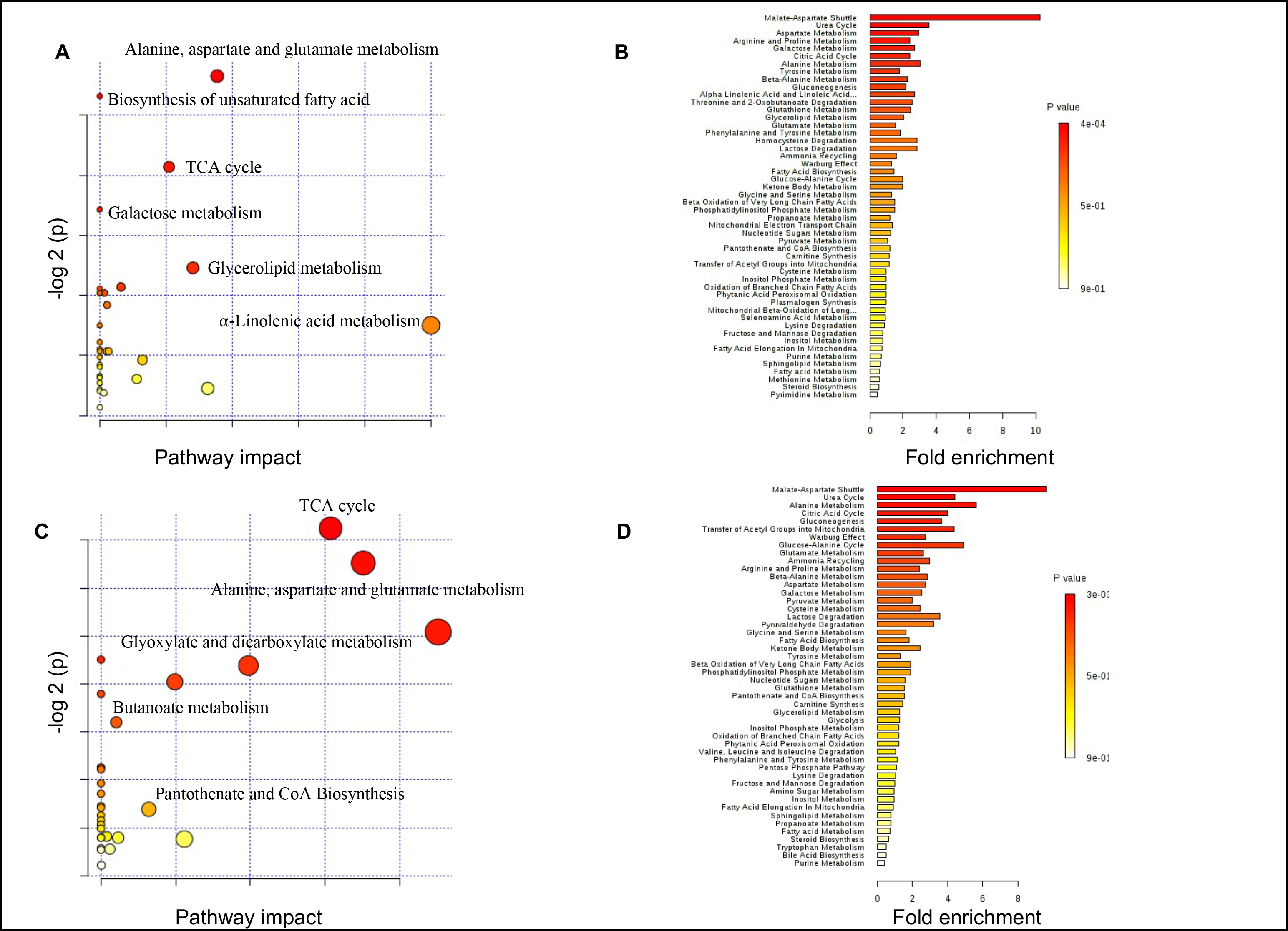

